# Mitochondrial cristae biogenesis coordinates with ETC complex IV assembly during *Drosophila* maturation

**DOI:** 10.1101/503474

**Authors:** Yi-fan Jiang, Hsiang-ling Lin, Li-jie Wang, Tian Hsu, Chi-yu Fu

## Abstract

Mitochondrial cristae contain electron transport chain complexes and are distinct from the inner boundary membrane (IBM) in both protein composition and function. While many details of mitochondrial membrane structure are known, the processes governing cristae biogenesis, including the organization of lipid membranes and assembly of nuclear and mitochondrial encoded proteins, remain obscure. We followed cristae biogenesis *in situ* upon *Drosophila* eclosion using serial-section electron tomography and revealed that the morphogenesis of lamellar cristae coordinates with ETC complex IV assembly. The membrane morphogenesis and functionalization were intricately co-evolved during cristae biogenesis. Marf-knockdown flies formed mitochondria of smaller sizes and reduced cristae content but organized lamellar cristae containing ATP synthase and functional COX. Instead, OPA1-knockdown flies had impaired cristae biogenesis and mitochondria function. We showed the ultrastructural localization of OPA1 in the cristae besides IBM that supports its functions in mediating cristae remodeling and inner membrane fusion. Overall, this study revealed the multilevel coordination of protein-coupled membrane morphogenesis in building functional cristae.

## Introduction

Mitochondria originate from an endosymbiosis event. The organelles exhibit unique double membrane architecture, consisting of outer and inner membranes that are separated by an intermembrane space. The inner membrane can be further subdivided into the inner boundary membrane (IBM) and the cristae invaginations, based on the ultrastructure, protein composition and function (Cogliati, Enriquez et al., 2016, Mannella, 2006). In the cristae, electron transport chain (ETC) complexes generate ATP by creating and maintaining a proton gradient between the matrix and intermembrane space (Gilkerson, Selker et al., 2003). Importantly, the morphology and remodeling of cristae are indicative of mitochondrial function, and cristae ultrastructure is known to be heavily influenced by several critical proteins (Barbot & Meinecke, 2016, Cogliati, Frezza et al., 2013, Frezza, Cipolat et al., 2006, Quintana-Cabrera, Mehrotra et al., 2018, Scorrano, Ashiya et al., 2002, Zick, Rabl et al., 2009). ATP synthase has been shown to play a structural role in inducing positive membrane curvature at the cristae ridges in addition to its enzymatic function (Davies, Strauss et al., 2011, Strauss, Hofhaus et al., 2008). Secondly, the mitochondrial contact site and cristae organizing system (MICOS) complex is known to stabilize the cristae junction, the region where cristae connect to the IBM (Huynen, Muhlmeister et al., 2016, Rampelt, Zerbes et al., 2017, Schorr & van der Laan, 2017). Optic atrophy protein 1 (OPA1), a protein involved in inner membrane fusion, also plays a pivotal role in stabilizing cristae junctions and mediating cristae remodeling during apoptosis (MacVicar & Langer, 2016, Varanita, Soriano et al., 2015). Even though key proteins have been identified as being essential for the maintenance and the remodeling of cristae architecture, the question of how functional cristae form *de novo,* including the organization of lipid membranes and assembly of proteins encoded by both nuclear and mitochondrial DNA, remains to be elucidated.

In this study, we investigated the mechanisms of cristae biogenesis by characterizing the mitochondrial development of *Drosophila* upon eclosion, the emergence of adult flies from pupa. At larval and pupal stages, *Drosophila* uses aerobic glycolysis to support the rapid growth of body mass and subsequent metamorphosis (Agrell, 1953, Tennessen, Baker et al., 2011). Their mitochondria contain only scarce lamellar cristae in the indirect flight muscle (IFM). Upon eclosion, mitochondria undergo development and establish densely arranged lamellar cristae that form connective membrane networks (Jiang, Lin et al., 2017b). It provides a well-characterized physiological time reference to study the *de novo* formation of functional cristae. Using this model system, we were able to uncover novel mechanisms of cristae biogenesis that involve intricate coordination of membrane morphogenesis and ETC complex IV assembly.

## Results

### Mitochondria exhibit cristae biogenesis upon *Drosophila* eclosion

To characterize mitochondrial structure in *Drosophila* during eclosion from the pupa, the IFM of adult flies was sampled at various time points. Thin-section TEM analysis showed that the mitochondria at day 1 after eclosion contained only a few organized cristae that were loosely scattered throughout the matrix (Fig 1a). The mitochondria developed densely packed lamellar cristae, usually within a couple days, to be what was observed in the matured flies (Fig 1b) (Jiang et al., 2017b). The mitochondrial protein abundance during maturation was analyzed. The mitochondria of day 1 flies contained a lower level of ATP5A (complex V of the ETC complex) compared to those of the week 4 flies according to the immuno-EM analysis (Fig 2a-b). The western blots of the whole fly extracts showed that at day 1, some other nuclear DNA-encoded mitochondrial proteins, such as pyruvate dehydrogenase (PDHA1), superoxide dismutase 2 (SOD2), and Cytochrome c (Cyt c), were approximately 30-50% of the levels in the week 4 flies (Fig 2c). On the other hand, the ribosomal protein, RPS6, was expressed roughly 18-fold more in the day 1 flies than week 4 flies (Fig 2c). Consistent with this observation, thin-section TEM micrographs revealed highly abundant ribosome or polyribosome-like structures in the cytoplasm of day 1 flies, but were much less abundant in the week 4 flies (Fig 1a-b). With this reliably orchestrated transition in mitochondrial morphology, the eclosion of *Drosophila* provides an excellent model system with which to elucidate the development of cristae ultrastructure and function *in situ.*

**Fig 1.**
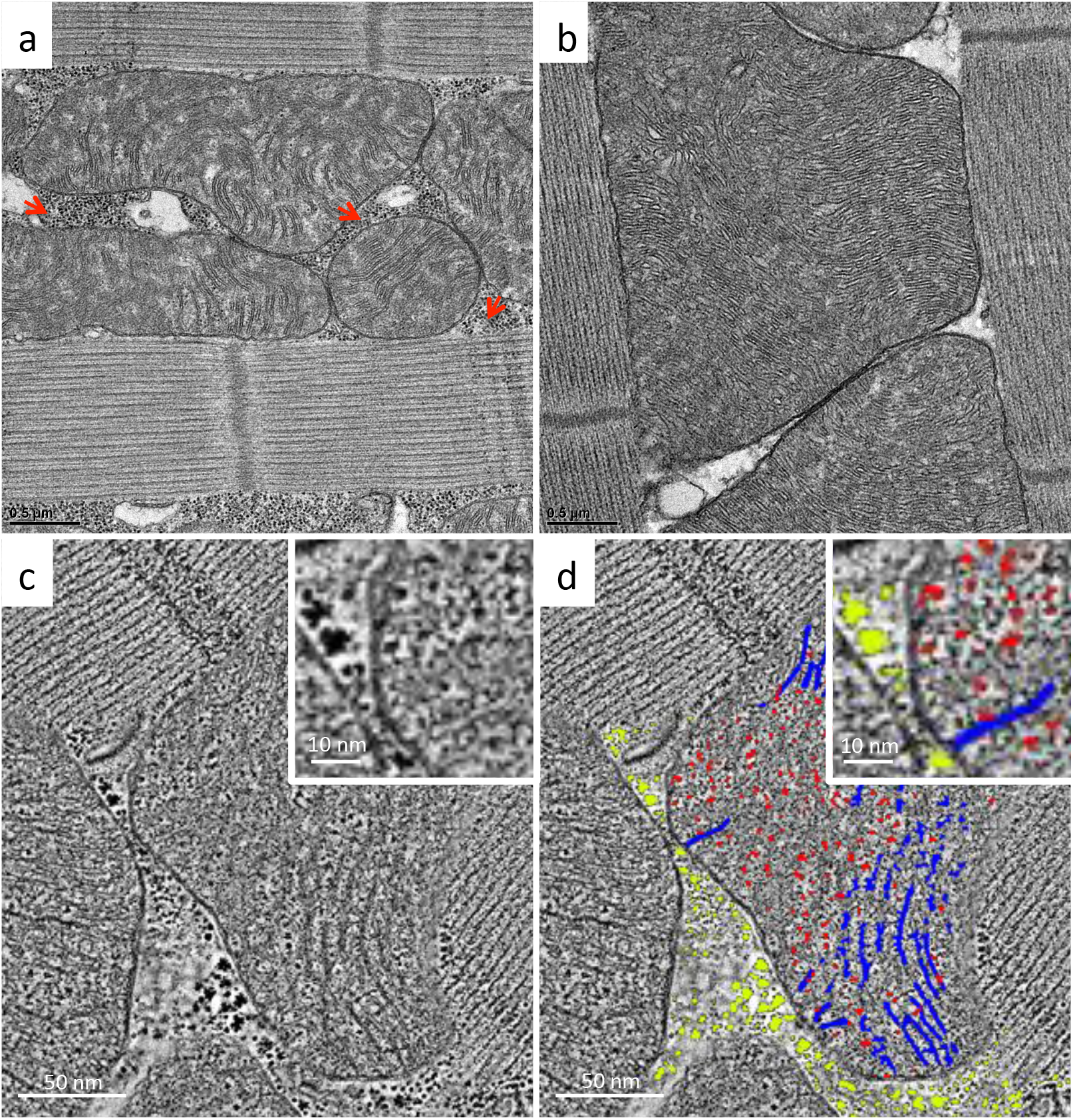
3D visualization of mitochondrial development upon *Drosophila* eclosion. A thin-section TEM micrograph of *Drosophila* IFM at day 1 (a) and week 4 (b). A slice of a serial-section tomography reconstruction (c) and the segmentation (d) of *Drosophila* IFM at day 1. (a) red arrows: cytoplasmic ribosomal-like densities; yellow arrows: close inter-mitochondrial contacts. (d) blue: cristae; red: mitochondrial ribosomal-like densities; green: cytoplasmic ribosomal-like densities.

**Fig 2.**
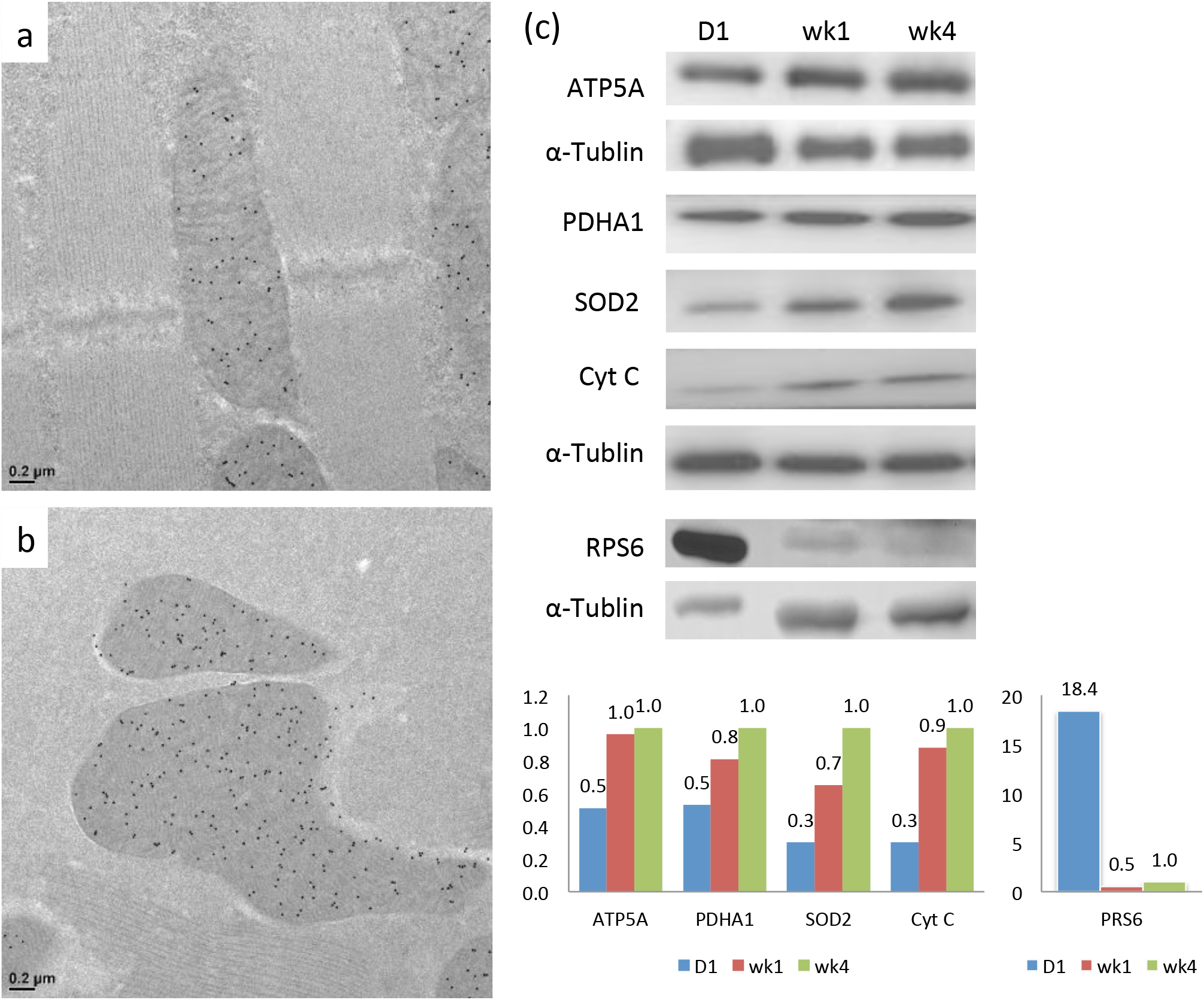
Analysis of protein contents during mitochondrial development. Immuno-EM labeling of ATP5A of *Drosophila* IFM at day 1 (a) and week 4 (b). Western-blot analysis of mitochondrial proteins ATP5A, PDHA1, SOD2, CytC and ribosomal protein RPS6 of *Drosophila* at day 1, week 1, and week 4 (c). The relative protein abundance was quantified by the densitometry and normalized to the signal of α-tubulin. The ratios were subsequently normalized to those of week 4.

To track cristae organization during mitochondrial maturation in 3D, we applied serial section electron tomography to reconstruct entire mitochondrial volumes. The global organization of IFM tissue was established upon eclosion with mitochondria distributed between parallel muscle fibers. In some cases, close inter-mitochondrial contacts were already observed, which may facilitate communication between mitochondria (Fig 1a, Movie 1) (Picard, McManus et al., 2015). Mitochondria in the day 1 flies appeared more polymorphic and contained lamellipodia-like or filopodia-like extensions, which became ovoid-shaped and filled the cytoplasmic space between the muscle fibers as they matured (Movie 1). Concentrated cytoplasmic ribosome or polyribosome-like densities surrounded the immature mitochondria, which would be expected to support rapid protein synthesis (Fig 1c-d). A cryo-tomography study reported that cytoplasmic ribosomes associate with the mitochondrial surface through the interaction with TOM complex (Gold, Chroscicki et al., 2017).

Joined serial tomograms of the day 1 flies showed that the mitochondrial matrix also contained numerous darkly stained ribosome-like molecules along with the segments of lamellar cristae (Fig 1c-d, Movie 1). In the mature mitochondria, mitochondrial ribosomes cannot be not readily identified in the densely confined matrix compartment. Due to the lack of available antibodies against *Drosophila* mitochondrial ribosome, the change of mitochondrial ribosome level during maturation was not quantified.

### Cristae biogenesis coordinates with COX complex assembly

Functional cristae require the proper organization of membrane and protein assembly. To investigate how membrane morphogenesis is coupled with functionalization, we took advantage of a traditional method of Cytochrome c oxidase (COX) staining to visualize COX activity in the context of the ultrastructure (Seligman, Karnovsky et al., 1968). COX oxidizes 3,3’-diaminobenzidine (DAB) and forms osmiophilic precipitants in the presence of osmium tetraoxide that appear darkly stained under TEM. The osmium tetraoxide substrate also binds to the head group of phospholipids that creates weak contrast for lipid membranes.

Mitochondria of the day 1 flies exhibited some lamellae of cristae with prominent COX activity, while some membranous structures that filled the matrix had weak staining (Fig 3a). To characterize their 3D arrangement, serial section electron tomography was applied. In the whole-mitochondria reconstructions, segments of lamellar cristae were observed scattered throughout the matrix (Fig 3c-e, Movie 2). The COX-negative membranes appeared as poorly organized reticulum. They contained very limited COX activity, therefore, we did not define them using the word “cristae”. The membranes gained COX activity as became organized as a part of the lamellae with a more defined width (Fig 3c-e, Movie 2).

**Fig 3.**
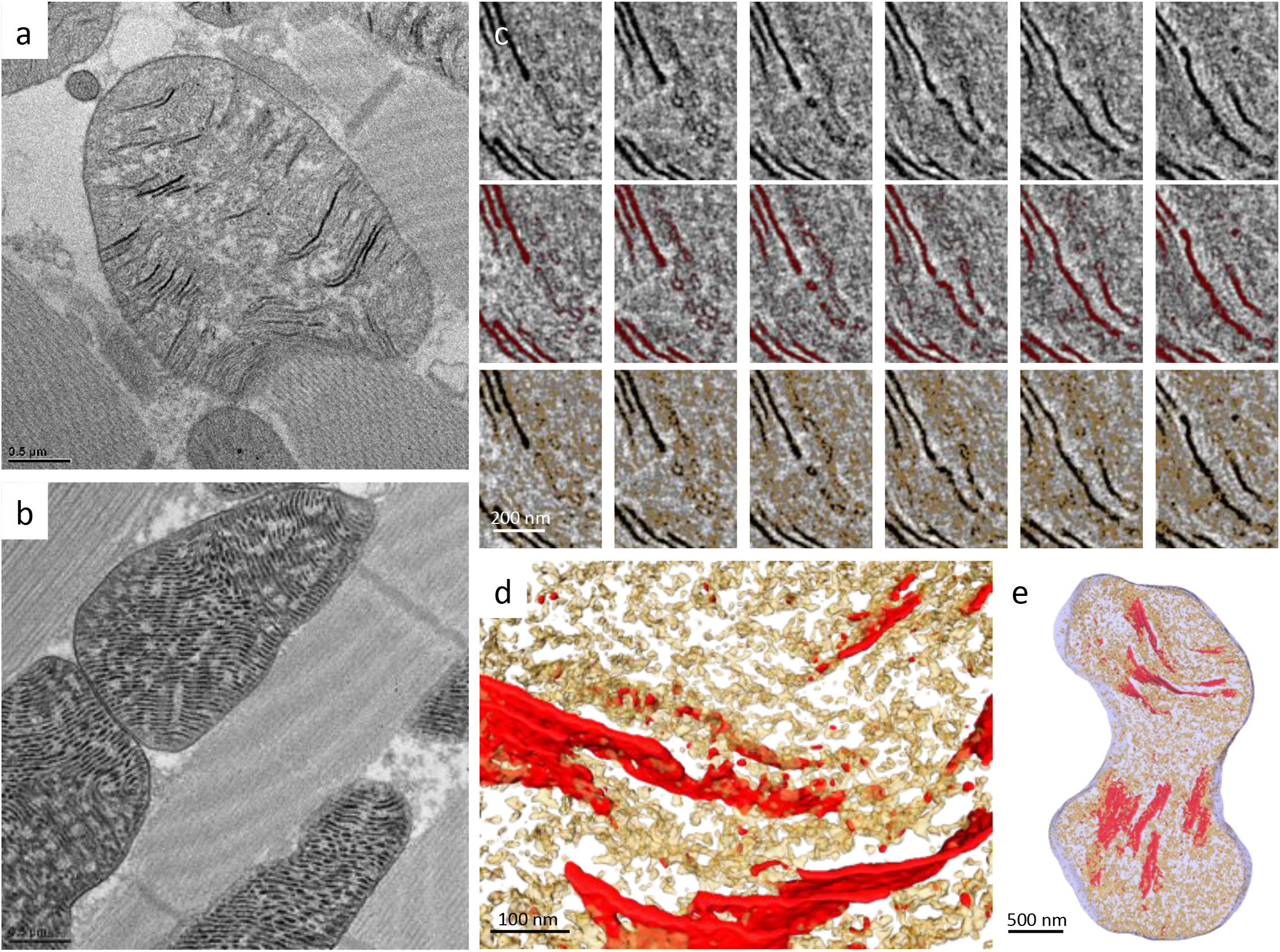
Cristae morphogenesis coordinates with COX assembly. A thin-section TEM micrograph of *Drosophila* IFM at day 1 (a) and week 4 (b) stained for COX activity. Tomographic slices across the z-axis and the corresponding segmentation of *Drosophila* IFM at day 1 stained for COX activity were shown (c). 3D representations of the tomographic segmentation were shown in (d) and (e). (c-e), red: COX-positive cristae; yellow: COX-negative reticular membranes. Positive COX activity appeared darkly stained in the micrographs. (a) red arrows: COX-positive cristae; yellow arrows: COX-negative membranes.

To verify the COX-staining results, we generated a knock-in fly that expresses Apex2 conjugated to the c-terminus of COX4, a subunit of COX that is synthesized in the cytoplasm and subsequently transported into the mitochondria (Fig S1a-b). Apex2, an ascorbate peroxidase, catalyzes the polymerization of DAB in the presence of hydrogen peroxide (H_2_O_2_), which enhances EM contrast after osmium tetraoxide staining, thus allowing us to track COX4 protein localization at the ultrastructural level (Martell, Deerinck et al., 2012). Using this method, COX4 was shown to localize mainly to the organized lamellar cristae in the day 1 flies, which correlated with the COX activity staining data (Fig 4a). The wild type flies were performed as the negative control of the Apex2 staining (Fig 4b). According to the structural studies, the c-terminus of COX4 is apposed to the intermembrane space, where the Apex2 staining appeared (Fig 4a) (Wu, Gu et al., 2016).

**Fig 4.**
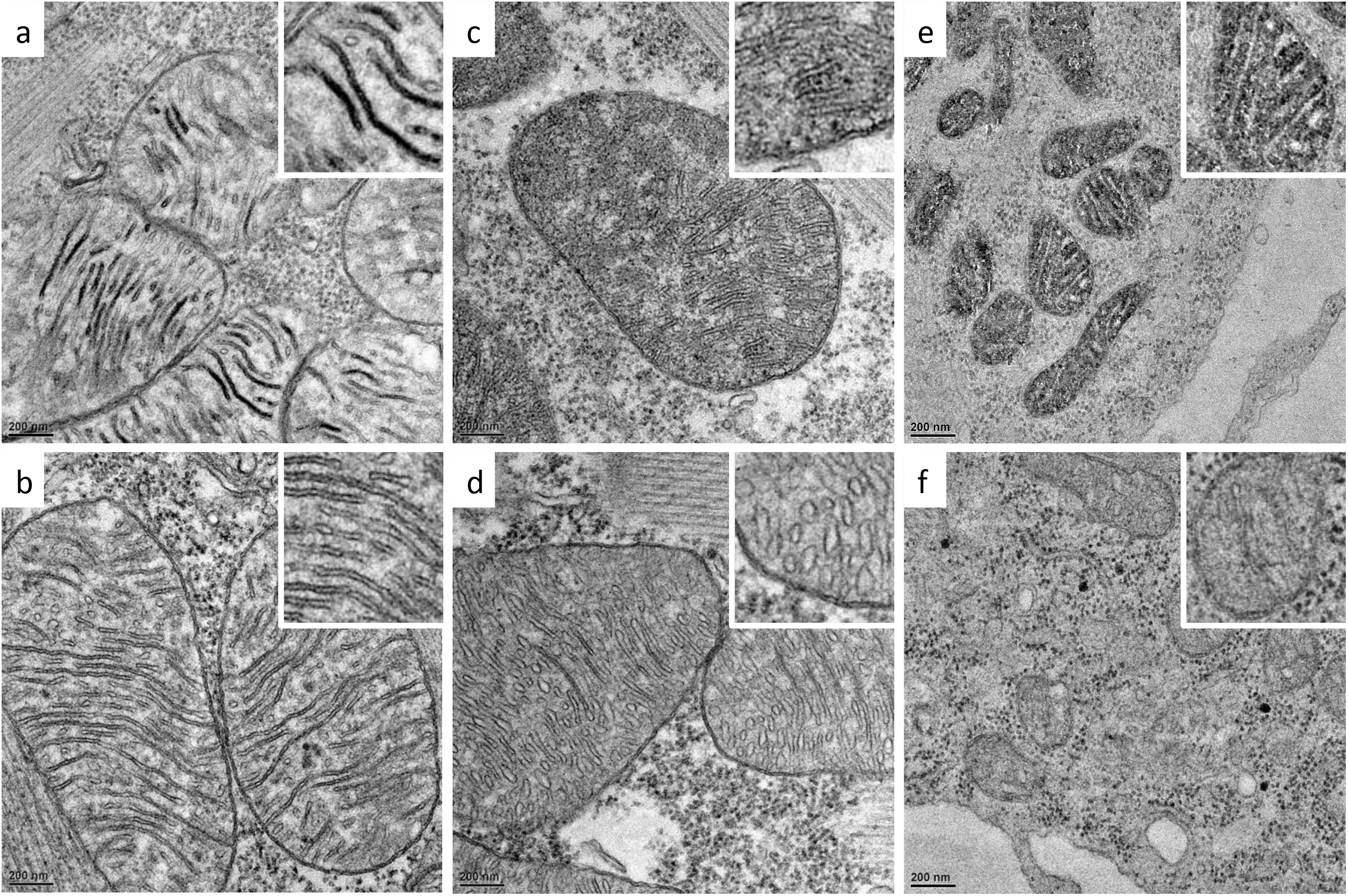
Ultrastructural tracking of COX4 and OXA1 during cristae biogenesis. Apex2 staining of the IFM of COX4-Apex2 knock-in flies (a) and OXA1-Apex2 knock-in flies (c) at day 1 were shown. Apex2 staining of the wild type at day 1 as negative controls were shown in (b) and (d), respectively. Apex2 staining of S2 cells transfected with and without plasmids expressing *D. melanogaster* OXA1-Apex2 was shown in (e) and (f), respectively. Positive Apex2 signals appeared darkly stained in the micrographs.

The COX complex comprises multiple subunits encoded by both nuclear and mitochondrial DNA. The insertion and assembly of the COX complex subunits require OXA1, which mediates the insertion of both nuclear and mitochondrial DNA-encoded polypeptides from the matrix into the inner membrane (Keil, Bareth et al., 2012, Soto, Fontanesi et al., 2012). OXA1 was shown to present in IBM and cristae in yeast by immuno-gold EM study (Stoldt, Wenzel et al., 2012). We set out to visualize if OXA1 also locates in the cristae to facilitate COX assembly during mitochondrial maturation in *Drosophila.* We tracked OXA1 localization in OXA1-Apex2 knock-in flies using the Apex2 method aiming for higher spatial resolution. Indeed Apex2 staining was present in the cristae and the IBM of the day 1 mitochondria (Fig 4c, S1a). The negative control of the Apex2 staining using the wild type flies was shown in Fig 4d. Judging by the staining location, the c-terminal Apex2 tag faced the matrix side of the inner membrane, and the staining pattern appeared as granular densities (Fig 4c). The result was confirmed in S2 cells that over-expressed OXA1-Apex2 (Fig 4e). The cells with mock-transfection were used as the negative control of Apex2 staining (Fig 4f). The western-blot of OXA1-Apex2 expression in S2 cells was shown in Fig S1c.

The data showed lamellar cristae are organized in coordination with the assembly of COX during cristae biogenesis upon *Drosophila* eclosion. The study revealed the intricate coordination of membrane morphogenesis and the acquisition of functionality.

### Marf-knockdown flies formed lamellar cristae containing ATP synthase and functional COX

Marf, a homolog of human mitofusin 1 and 2, mediates outer membrane fusion and influences ER-mitochondria tethering (de Brito & Scorrano, 2008, Detmer & Chan, 2007, Filadi, Greotti et al., 2015, Filadi, Pendin et al., 2018, Schrepfer & Scorrano, 2016). To investigate how Marf affects cristae biogenesis and mitochondrial maturation, we investigated the mitochondrial structure and function of Marf-knockdown flies. Marf-knockdown flies had compromised climbing ability (Fig S2a). The whole fly extracts had similar levels of ATP5A, PDHA1, SOD2, and Cyt c as compared to the wild type flies (Fig S2b). Thin-section TEM and serial-section tomography revealed that lamellar cristae were formed in Marf-knockdown mitochondria (Fig 5a-d, Fig S2c-d, Movie 3). In addition, they contained COX activity (Fig 5e, S2d), as well as ATP5A and Cyt c shown by the immuno-EM (Fig 5f, Fig S2f). The Apex2 staining of the ATP synthase OSCP-Apex2 knock-in flies under the Marf-knockdown background also confirmed the presence of ATP synthase OSCP in the cristae (Fig 5g, Fig S2h). The dark staining of OSCP-Apex2 was restricted to the matrix side of the cristae. It correlated with the structural studies of properly assembled ATP synthase and suggested correct folding and targeting of OSCP-Apex2 in the knock-in flies (Fig 5g) (Wu et al., 2016, Zhou, Rohou et al., 2015).

**Fig 5.**
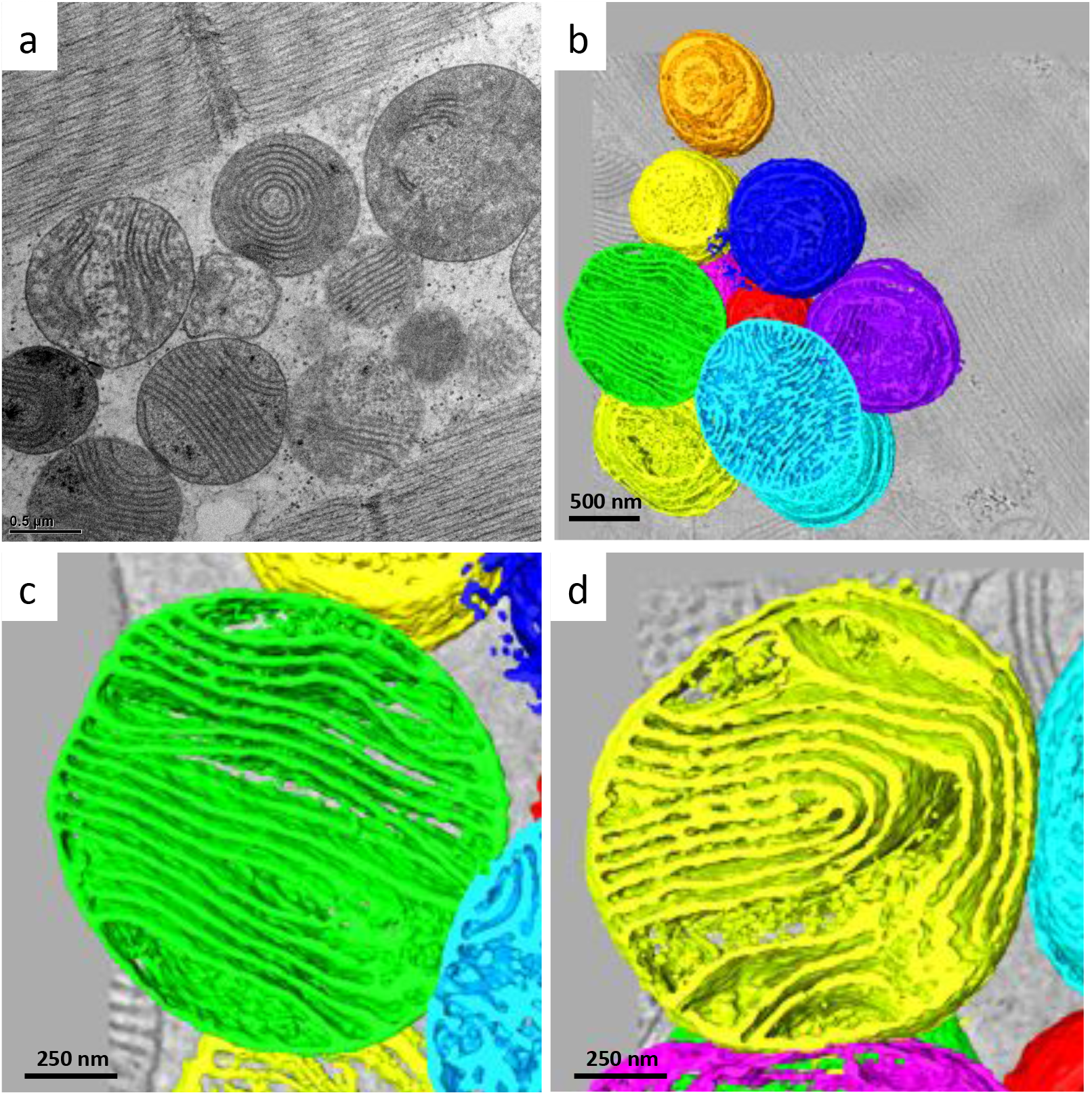
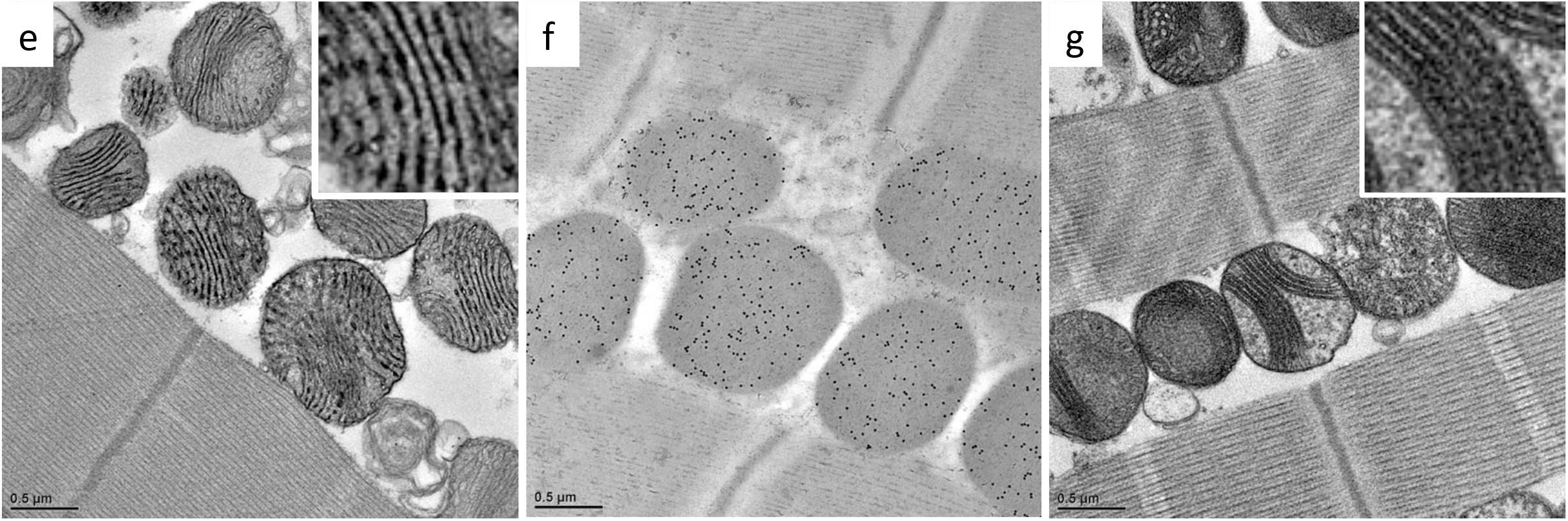
Marf-knockdown flies formed lamellar cristae containing COX and ATP synthase. Thin-section TEM micrograph (a), the tomographic segmentation (b-d), COX activity staining (e), immuno-EM against ATP5A (f), and ATP synthase OSCP-Apex2 staining (g) of Marf-knockdown flies at week 4 were shown. Positive Apex2 signal or COX activity appeared darkly stained in the micrographs.

Even though Marf-knockdown mitochondria organized lamellar cristae that contain COX and ATP synthase, they were approximately 49% smaller than the wild type, which was expected given that mitochondrial outer membrane fusion was impaired. In addition, the cristae content per mitochondrial volume was reduced to approximately 53% in the Marf-knockdown flies, comparing to approximately 99% in the wild type at week 4. This observation probably reflects the alteration of ER-mitochondria contacts in the Marf-knockdown, which is essential for lipid transport to the mitochondria from the ER (Area-Gomez, Del Carmen Lara Castillo et al., 2012, Filadi et al., 2018, Tatsuta & Langer, 2017, Vance, 2014).

### OPA1-knockdown flies showed impaired cristae biogenesis and function

We investigated how OPA1 affects cristae biogenesis and function. OPA1-knockdown flies showed reduced climbing ability (Fig S3a). Several nuclear DNA-encoded mitochondrial proteins in the whole fly extracts were at a similar level as in the wild type flies (Fig S3b). OPA1-knockdown mitochondria were approximately 42 % smaller than the wild type mitochondria, as a result of defective mitochondrial fusion. Many mitochondria contained very few organized membranes but vacuoles, some likely resulting from the incomplete inner membrane fusion post the outer membrane fusion (Fig 6a, Fig S3c). Most OPA1-knockdown mitochondria had disordered and aberrant membranes.

**Fig 6.**
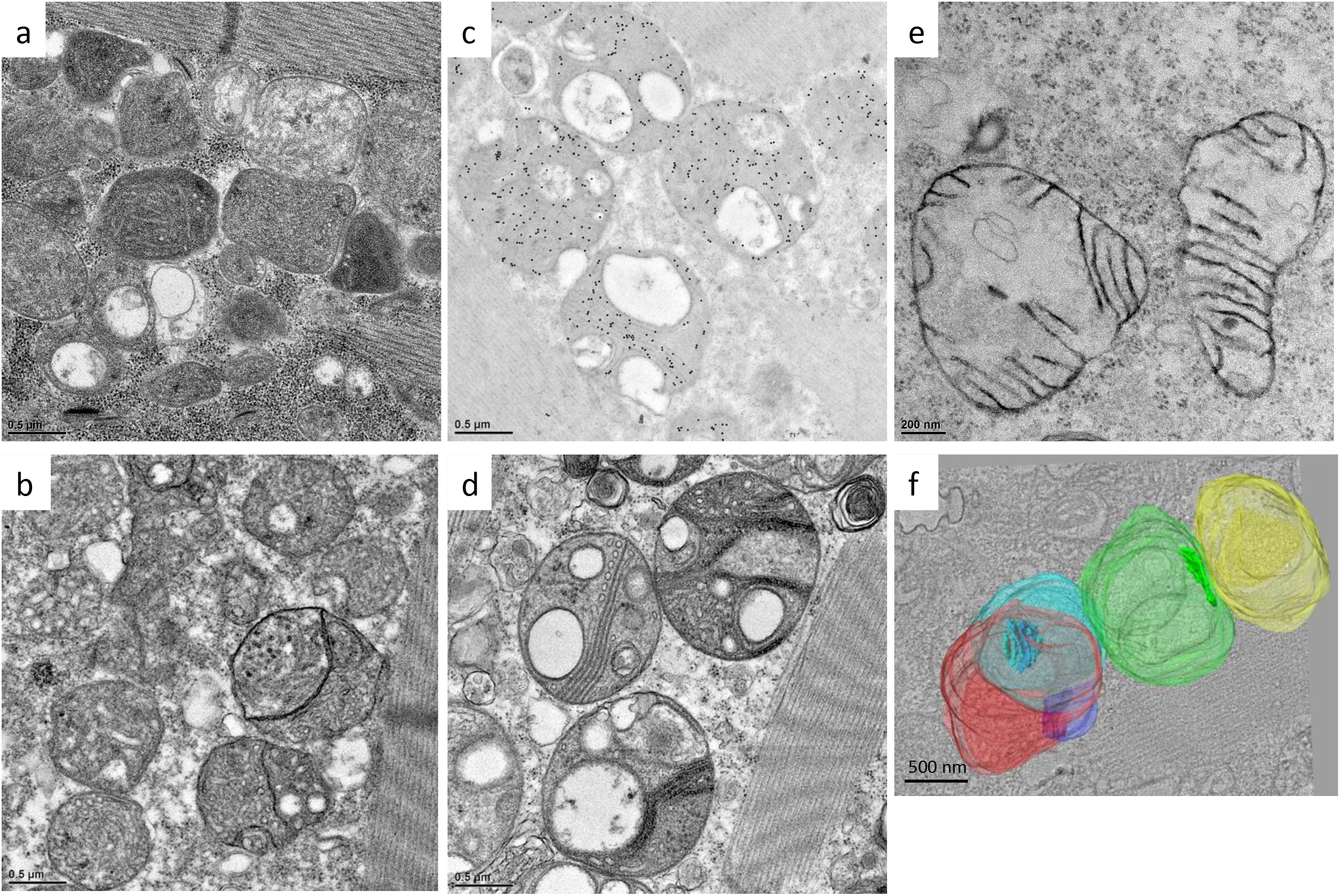
OPA1-knockdown flies showed impaired cristae biogenesis and function. OPA1-knockdown flies at day 1 were analyzed by thin-section TEM (a), COX activity staining (b), immuno-EM against ATP5A (c), ATP synthase OSCP-Apex2 staining (d), and the tomographic segmentation (f). Sub-mitochondrial localization of OPA1 was tracked in 293T cells overexpressing human OPA1-Apex2 (e).

OPA1-knockdown mitochondria also had very low levels of functional COX assemblies in both day 1 and week 4 flies (Fig 6b, Fig S3d, Movie 4). Only about 5% of the mitochondria exhibited positive COX staining. Immuno-EM suggested a reduced level of ATP5A (F_1_ subunit α) and cytochrome c proteins in OPA1-knockdown mitochondria (Fig 6c, Fig S3e-g). In correlation, ATP synthase OSCP-Apex2 knock-in conjugates under OPA1-knockdown background only appeared in a few regional lamellar membranes (Fig 6d). Dysmorphic cristae ultrastructure and decreased ETC assemblies likely led to a vicious circle of low membrane potential and impaired protein transport (Harbauer, Zahedi et al., 2014, Song, Chen et al., 2007). OPA1 was also shown to affect mitochondrial DNA maintenance that likely contributed to the poor mitochondrial function and morphogenesis (Elachouri, Vidoni et al., 2011).

We tracked OPA1 localization by Apex2 conjugation in 293T cells overexpressing human OPA1-Apex2 (Fig 6e, S4a-b), as well as in S2 cells overexpressing *D. melanogaster* OPA1-Apex2 (Fig S4c-d). OPA1 protein was observed in both cristae and the IBM. Opa1-Apex2 knock-in flies had very low expression of OPA1-Apex2, approximately 7% of the expression of COX4-Apex2 knock-in by western blot against Flag tag (Fig S1a, S4e-f). The ultrastructural localization of OPA1 supports its roles in mediating inner membrane fusion and cristae remodeling.

## Discussion

In this study, we took the advantage of *Drosophila* model that displayed dramatic cristae biogenesis upon eclosion in building compact lamellar cristae. We showed that the development of cristae morphology and functionality is intricately coordinated. The COX complex is composed of subunits encoded by both nuclear and mitochondrial DNA. The assembly pathway has been described in great detail and involves the coordination of multiple steps, including protein synthesis, membrane insertion, assembly, and metal incorporation, all of which are mediated by various chaperones and accessory proteins (Soto et al., 2012). Our study uncovered an extra layer of coordination between the assembly of COX and the establishment of initial cristae ultrastructure. Previously, ATP synthase has been demonstrated essential in cristae morphogenesis (Davies et al., 2011, Strauss et al., 2008), so as cristae morphology to determine the ETC supercomplex assembly and respiratory efficiency (Cogliati et al., 2013). The intimate connection of cristae morphology with ETC assembly and function is reiterated in this study.

Marf is a homolog of human mitofusin 1 and mitofusin 2. It is involved in mitochondrial outer membrane fusion, ER-mitochondria contact, and neuromuscular function (de Brito & Scorrano, 2008, Detmer & Chan, 2007, Filadi et al., 2015, Filadi et al., 2018, Khalil, Cabirol-Pol et al., 2017, Sandoval, Yao et al., 2014, Schrepfer & Scorrano, 2016). Marf-knockdown flies display smaller size of mitochondria as a phenotype of inhibiting outer membrane fusion. They also have reduced cristae membrane content. Most mitochondrial lipids or the precursors are imported from ER through the proteins mediating membrane contacts and lipid translocation (Flis & Daum, 2013, Tatsuta & Langer, 2017). Cardiolipin, in particular, was shown to stabilize ETC components and shape cristae architecture (Desmurs, Foti et al., 2015, Paradies, Paradies et al., 2014). Significant reduction of cristae content per mitochondrial volume was observed in the Marf-knockdown. However, cristae biogenesis is not noticeably affected as they form lamellar cristae harboring ATP synthase and functional COX. Previous studies also showed normal cristae morphology and ETC supercomplex assembly, even though mtDNA copy number is reduced (Cogliati et al., 2013).

On the contrary, OPA1-knockdown flies display severe impairment in cristae morphogenesis and function, likely reflects its essential role in various aspects of mitochondrial processes. OPA1, a dynamin-related GTPase, is anchored in the inner membrane in its long isoform. Upon proteolytic cleavage, the short isoform becomes localized to the intermembrane space. This proteolysis-mediated redistribution is an important regulatory mechanism in OPA1-mediated membrane fusion and fission (Ban, Ishihara et al., 2017, Del Dotto, Mishra et al., 2017, MacVicar & Langer, 2016). OPA1 has also been shown to control cristae remodeling and junction widening during apoptosis (Frezza et al., 2006). It interacts with MICOS component Mic60 in the cristae and cristae junction (Hoppins, Collins et al., 2011, Sastri, Darshi et al., 2017). OPA1 protects mitochondria from complex III inhibition by stabilizing cristae morphology and ATP synthase oligomers (Quintana-Cabrera, Quirin et al., 2018). In this study, we reported the ultrastructural localization of OPA1 in the cristae besides the IBM, which supports its multiple roles in cristae remodeling and inner membrane fusion.

Through evolution, mitochondria have been delicately integrated as organelles, which contain highly functionalized compartments and membranes. Thus, it is no surprise that a sophisticated biomolecular interaction network is being uncovered in the regulation of cristae architecture (Jayashankar, Mueller et al., 2016). A model of cristae formation was proposed previously where one pathway involves mitochondrial fusion and OPA1-mediated inner membrane fusion, while in another cristae are formed through the invagination of IBM independent of mitochondrial fusion (Harner, Unger et al., 2016). In the study, we demonstrated the intricate multilevel coordination of building functional cristae. The generalized mechanism of protein-coupled membrane morphogenesis was clearly demonstrated for cristae biogenesis, which is in line with various other membrane remodeling processes, such as vesicle budding and fusion (Bonifacino & Glick, 2004).

## Materials and Methods

### Fly strains

*Drosophila* strains on the Oregon-R-P2 background were used in these studies. Marf-knockdown flies were obtained by crossing UAS (Bloomington 31157) and GAL4 (Bloomington 26882) lines. OPA1-knockdown flies were obtained by crossing UAS (Bloomington 32358) and GAL4 (Bloomington 38459) lines.

Apex2-Flag knock-in flies of COX4, OXA1, ATP synthase OSCP, and OPA1 were generated by CRISPR/Cas9-mediated genome editing and homology-dependent repair using a guide RNA(s) and a dsDNA plasmid donor. PBac system was used to facilitate genetic screening (Well Genetics). To generate ATP synthase OSCP-Apex2 under Marf-knockdown background, a stable line was selected by crossing ATP synthase OSCP-Apex2 line with GAL4 (Bloomington 26882) line, which was subsequently crossed with UAS (Bloomington 31157) line. To generate ATP synthase OSCP-Apex2 under OPA1-knockdown background, a stable line was selected by crossing ATP synthase OSCP-Apex2 line with GAL4 (Bloomington 38459) line, which was subsequently crossed with UAS (Bloomington 32358) line.

### HPF/FS specimen preparation for morphological observation

Flies were anesthetized on ice and embedded in 4% low melting agarose in 0.1 M phosphate buffer. Embedded flies were then sectioned to 100 μm-thick slices by a vibrating blade microtome (Leica VT1200S) and fixed in 2.5% glutaraldehyde in phosphate buffer.

HPF/FS was also performed as previously described (Jiang, Lin et al., 2017a, Jiang et al., 2017b). The tissue sections were washed in 3 drops (~150 μl) of phosphate buffer, followed by 2 drops (~100 μl) of phosphate buffer with 20 *%* BSA. The specimens were subsequently placed in the gold carrier filled with 20 % BSA in PBS. The carriers were loaded into a high-pressure freezer (Leica EM HPM100) according to manufacturer’s instructions. The carriers were subsequently released from the holder under liquid nitrogen and transferred to the chamber of a freeze-substitution device (Leica EM AFS2) pre-cooled to −140 °C and incubated for 96 hr before FS.

During FS, the temperature of the chamber was raised to 0 °C at a slope of 5 °C/hr. During the process, the specimens were substituted with 0.1 % uranyl acetate and 2 % glutaraldehyde in acetone at −60 °C for 12 hr, followed by 2 % osmium tetroxide at −25 °C for 12 hr, and washed with acetone at 0 °C three times for 1 hr each. The specimens were subsequently removed from the carriers using a needle, infiltrated and embedded in EMBed-812 resin at room temperature, which was polymerized at 65 °C for 16 hr. The specimen blocks were trimmed and sectioned using an ultramicrotome. The sections were stained with Reynold’s lead citrate for 10 min and subjected to TEM inspection.

### Serial-section electron tomography

The procedure was also performed as previously described (Jiang et al., 2017a, Jiang et al., 2017b). Serial sections with a thickness of 200 nm were prepared and collected on copper slot grids (2 x 0.5mm oval slots) with carbon supports, on which overlaid with 10 nm fiducial gold pretreated with BSA. The grids were stained with Reynold’s lead citrate before the second layer of fiducial gold was applied. The specimens were imaged with FEI Tecnai TEM operating at 200 kV and the micrographs were recorded with a Gatan UltraScan 1000 CCD at 0.87 nm/pixel (9,600x). Tilt series from −60° to +60° with 2° increments were acquired at 10 μm defocus using Leginon automatic data collection software (Suloway, Shi et al., 2009). Double tilt series were collected using a double tilt holder (Model 2040 Dual-Axis Tomography Holder, Fischione). Serial tomograms were reconstructed, joined using IMOD, and segmented using Avizo 3D software (FEI).

### EM staining for COX activity

The procedure was modified as previously described (Seligman et al., 1968). Vibrating blade microtome sections of the fly tissues were washed with PBS and stained for 3 hr at 37 °C in a staining solution that contained 5 mg 3,3’-diaminobenzidine tetrahydrochloride (DAB), 9 ml sodium phosphate buffer (0.05M, pH7.4), 750 mg sucrose, 20 μg catalase (dissolved in 0.05M potassium phosphate buffer, pH 7.0), and 10 mg cytochrome c (dissolved in distilled water) at a volume of 10 ml. Subsequently, the specimens were washed with PBS for 1 hr and subjected to standard osmium fixation, dehydration, infiltration and embedded using Embed-812 resin. The blocks were cut to thin-sections of 70 nm thicknesses and observed under TEM without further staining.

### HPF/FS specimen preparation for immuno-EM labeling

Flies were sectioned in fixatives containing 4 % paraformaldehyde, 0.25 % glutaraldehyde in phosphate buffer and subjected to HPF/FS as described above with some modifications. Immuno-EM specimens were freeze-substituted with 0.1 % uranyl acetate in acetone at −90 °C for 58 hr (agitated every 8 hr), and warmed up to −45 °C at a slope of 5 °C /hr, washed with acetone three times for 1 hr each. The specimens were subsequently infiltrated through an ascending gradient of Lowicryl HM20 resin (10%, 20%, 40%, 60%, 80% and 90%, 8hr for each concentration, and agitated every 2 hr). The chamber was further warmed up to −25 °C at 5 °C /hr. The solutions were replaced with 100 % HM20 three times for 24 hr each (agitated every 2 hr). After adjusting the orientation within the carriers, ultraviolet polymerization was performed at −25 °C for 48 hr. The chamber was later warmed up to 20 °C (5°C /hr) and exposed to ultraviolet radiation for another 48 hr.

After polymerization, the specimen blocks were detached from HPF carriers. 100 nm thick sections were prepared and placed on 200-mesh nickel grids for immuno-EM labeling.

### Immuno-EM labeling

Thin sections placed on nickel grids were blocked with 10 *%* BSA in PBS for 20 min and incubated with primary antibodies in incubation buffer (1% BSA in PBS) for 2 hr. Grids were subsequently washed with incubation buffer three times (10 min each). Secondary antibodies, goat anti-mouse IgG (EM.GMHL15, BB International) and protein A (EM.PAG15, BB International) conjugated to 15 nm gold particles, were used against the primary antibodies from mouse and rabbit respectively. Secondary antibodies at 20-fold dilution were applied and samples were incubated for 1 hr. After washing with PBS, the immune-complexes were fixed with 1% glutaraldehyde in PBS and washed three times with distilled water. The specimens were inspected by TEM operating at 120 kV (FEI Tecnai G2 TF20 Super TWIN).

The primary antibodies and applied dilution factors are listed as follows: mouse anti-dsDNA (500x, abcam ab27156), mouse anti-ATP5A (500x, abcam ab14748), mouse anti-Cytochrome C (8000x, abcam ab13575), mouse anti-PDHA1 (500x, abcam ab110334), mouse anti-Ubiquitin (1000x, abcam ab7254), mouse anti-DNA-RNA hybrid (500x, kerafast ENH001), and rabbit anti-SOD2 (500x, abcam ab13534).

### Apex2 staining EM

The protocol was modified as previously described (Hung, Udeshi et al., 2016). Vibratome sections of the fly tissues were fixed in 2% glutaraldehyde in 0.1 M sodium cacodylate with 2 mM CaCl_2_, pH 7. Residual glutaraldehyde was quenched with 20 mM glycine followed by the washing steps. The specimens were subsequently stained with 0.5 mg/ml DAB-4HCL (3,3’-diaminobenzidine) and 0.3% H_2_O_2_ in the buffer for 30 min, washed, and stained with 1% osmium tetroxide for 30 min. After washes, the specimens were stained with 1% uranyl acetate overnight. The specimens were further dehydrated and embedded in resin for thin-section and TEM observation.

### Cell culture for Apex2 staining

293T cells were seeded on plastic membranes in a 6-well culture plate. The cells reached >80% confluence after overnight culture and were transfected with pECFP-OPA1 (human isoform1)-Apex2-Flag using *Trans*IT-X2 (Thermo-Fisher). After incubation overnight, the monolayer cells were fixed with 2% glutaraldehyde and followed the Apex2 staining procedure as described above (Hung et al., 2016).

S2 cells were seeded in a 6-well culture plate at 1*10^6^ cells/ml and grew for another day to 2-4*10^6^ cells/ml. The cells were transfected with pMT-V5-HisB-OXA1 (*D. melanogaster*)-Apex2-Flag or pMT-V5-HisB-OPA1 (*D. melanogaster*)-Apex2-Flag using calcium phosphate transfection kit (Invitrogen) and induced protein expression by CuSO_4_. The cells were harvested 2-3 days post-induction, fixed with 2% glutaraldehyde and followed the Apex2 staining procedure as described above (Hung et al., 2016).

### Western blot analysis

Flies were dissected and homogenized by Dounce tissue grinder in RIPA buffer containing protease inhibitor (cOmplete™, Roche). Cellular debris was removed by centrifugation at 4°C, 14000 x g for 20 min. The supernatants were collected and the protein concentrations were determined by Pierce protein assay (Pierce 660 nm Protein Assay Reagent, ThermoScientific). 0.6 μg/well of proteins were loaded for SDS-PAGE and western-blot analysis.

The antibodies used in the studies were as follows: mouse anti-ATP5A (50000x, abcam ab14748), mouse anti-Cytochrome C (10000x, abcam ab13575), mouse anti-PDHA1 (1000x, abcam ab110334) or rabbit anti-SOD2 (10000x, abcam ab13534), and rabbit anti-alpha tubulin (10000x, abcam ab18251), anti-mouse IgG-HRP (2000x, Invitrogen 62-6520) or anti-rabbit IgG-HRP (5000x, abcam ab97051). For quantification, ratios of the densitometry signal of individual proteins to that of alpha-tubulin were calculated. The ratios were then normalized to the wild type at week 4.

### Mitochondria analysis

The changes in the mitochondrial size of Marf and OPA1-knockdown vs. the wild type were calculated using thin-section EM micrographs. For each type, over a hundred mitochondria were analyzed. The cristae to mitochondrial membrane content of Marf-knockdown and the wild type 3D tomograms were analyzed using the automatic segmentation and analysis of Avizo 3D.

### Climbing assay

Flies were knocked down to the bottom of the culture tubes. Numbers of flies climbing over the target line (about 18cm) over 3 min were recorded. About 15 flies were used for each triplicate.

## Acknowledgments

We thank the EM core of Institute of Cellular and Organismic Biology and the cryo-EM core of Academia Sinica, Taiwan. We thank Dr. Ya-Hui Chou for the helpful discussion on *Drosophila* genetics. We thank the funding support from Academia Sinica AS-105-TP-B04 and MOST 105-2628-B-001-004-MY3.

## Author contributions

Yi-fan Jiang and Chi-yu Fu designed the experiments; Yi-fan Jiang, Hsiang-ling Lin, Li-jie Wang, and Tian Hsu performed the experiments; Chi-yu Fu wrote the paper.

---

**Fig S1.**
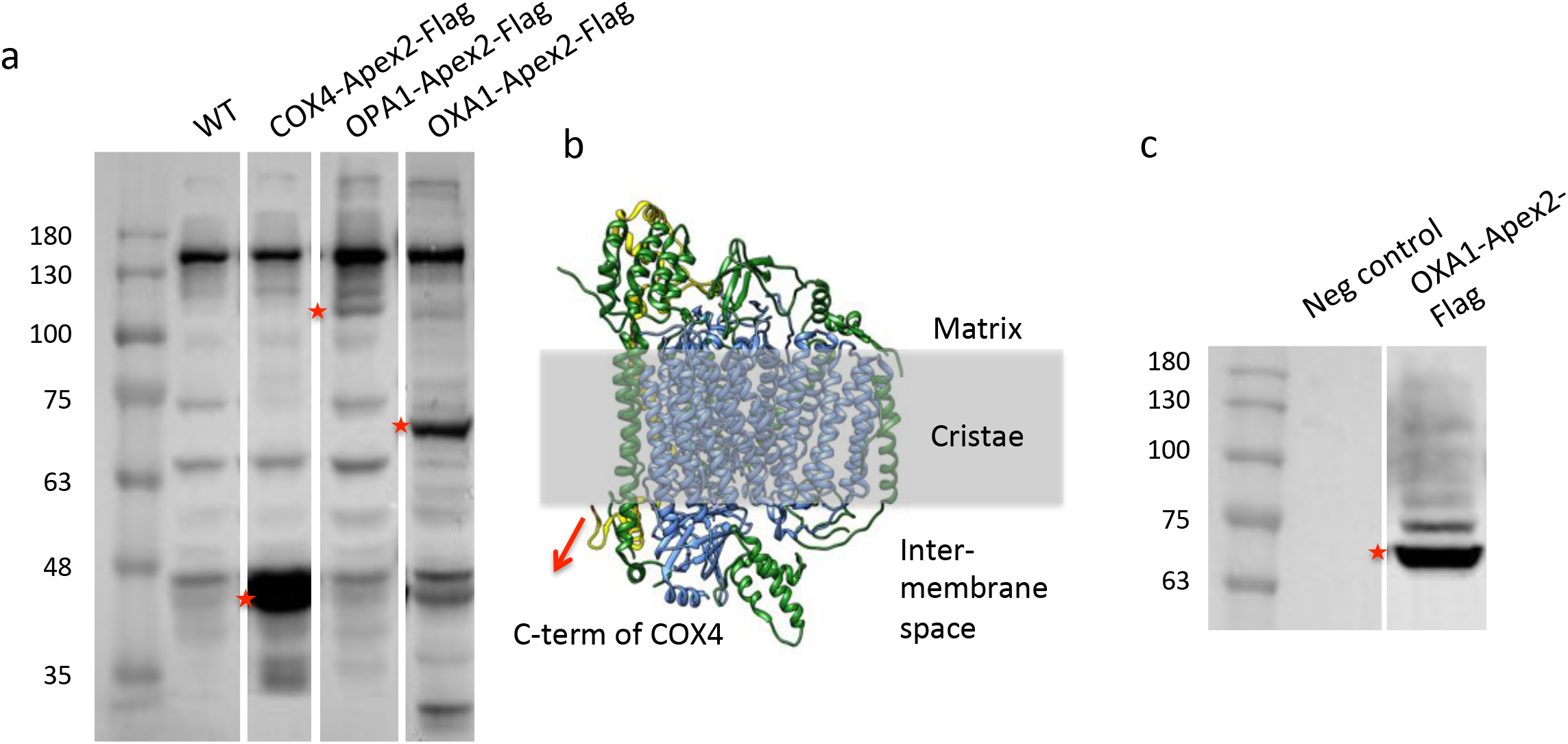

**Fig S2.**
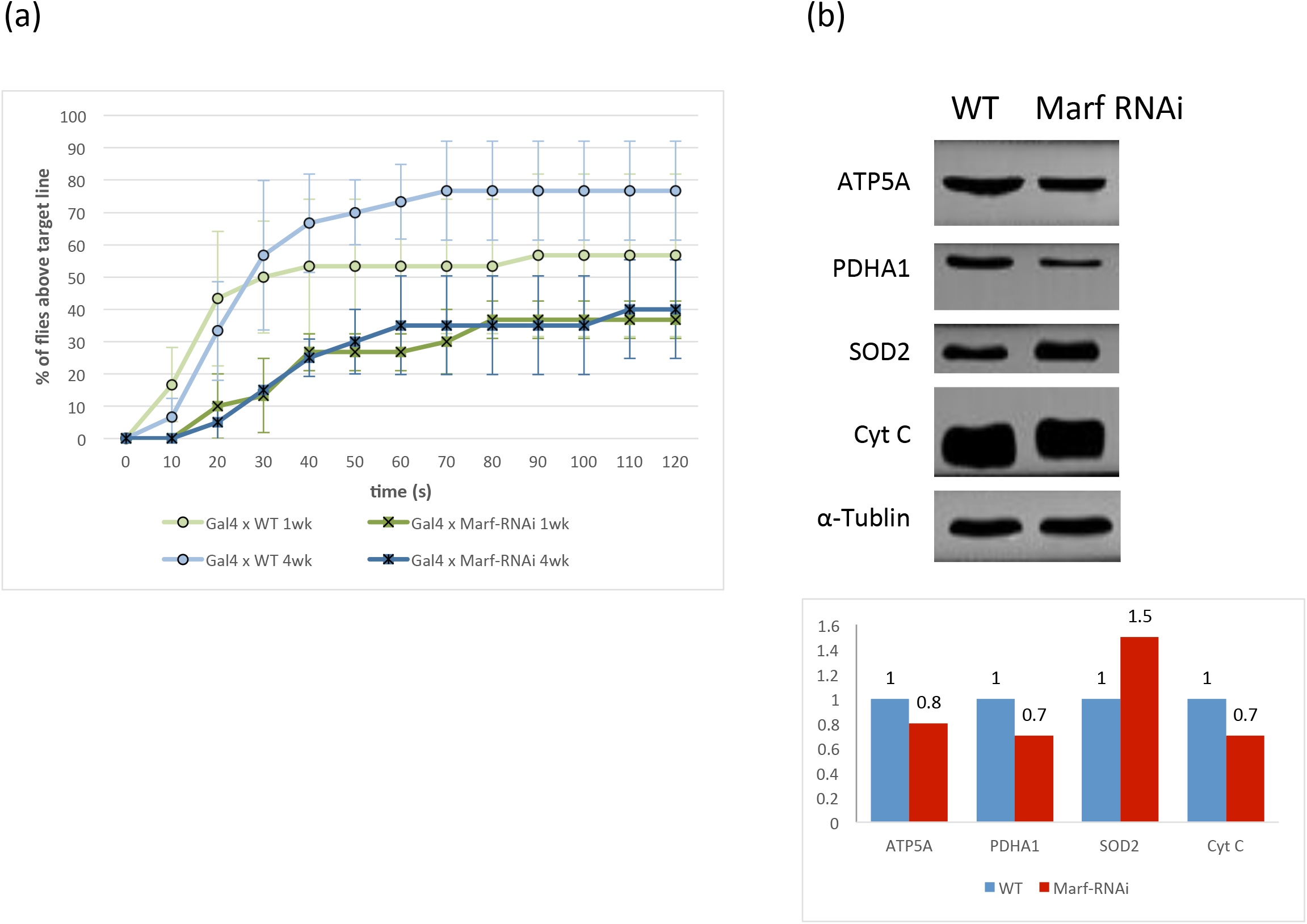
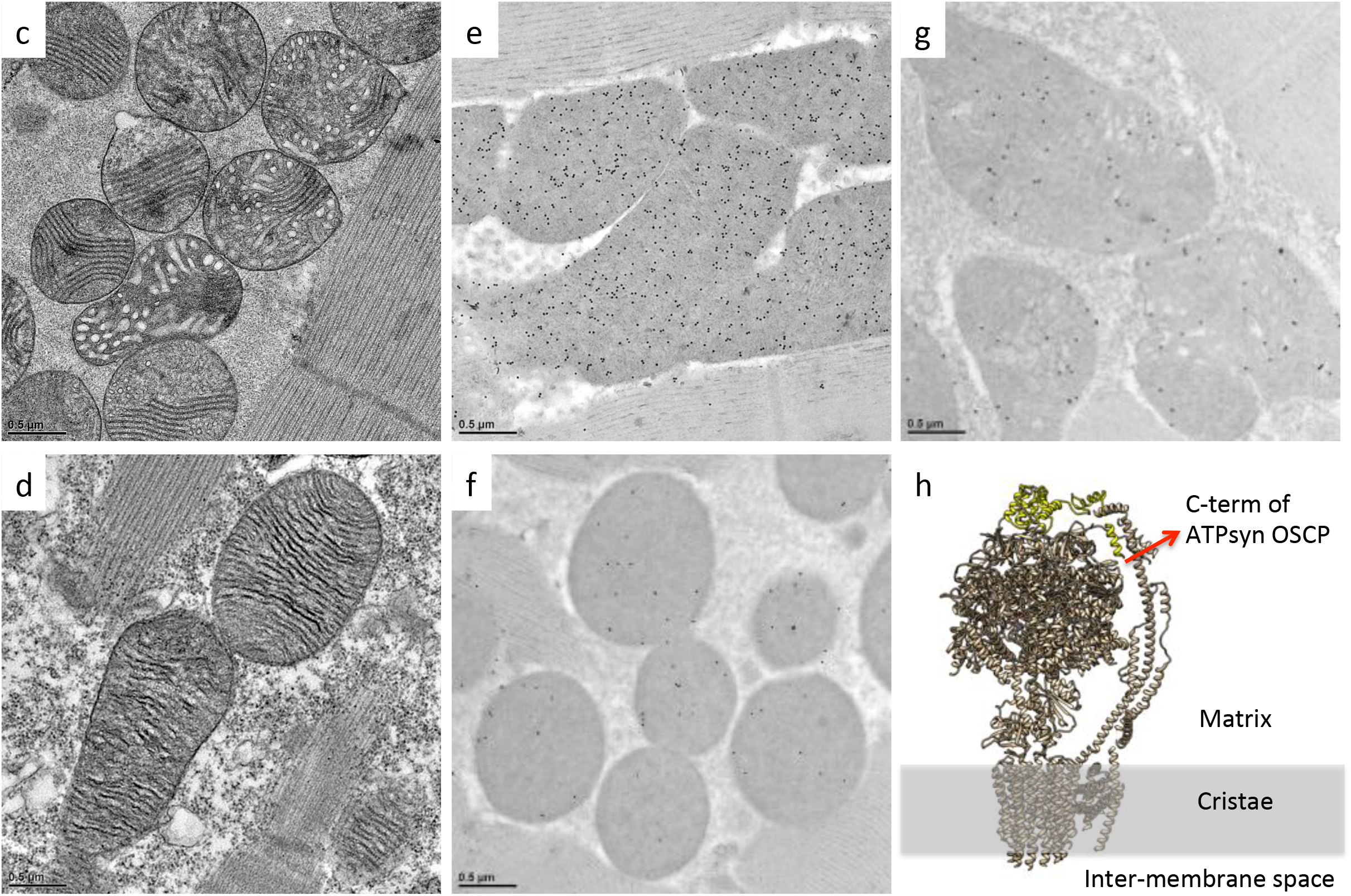

**Fig S3.**
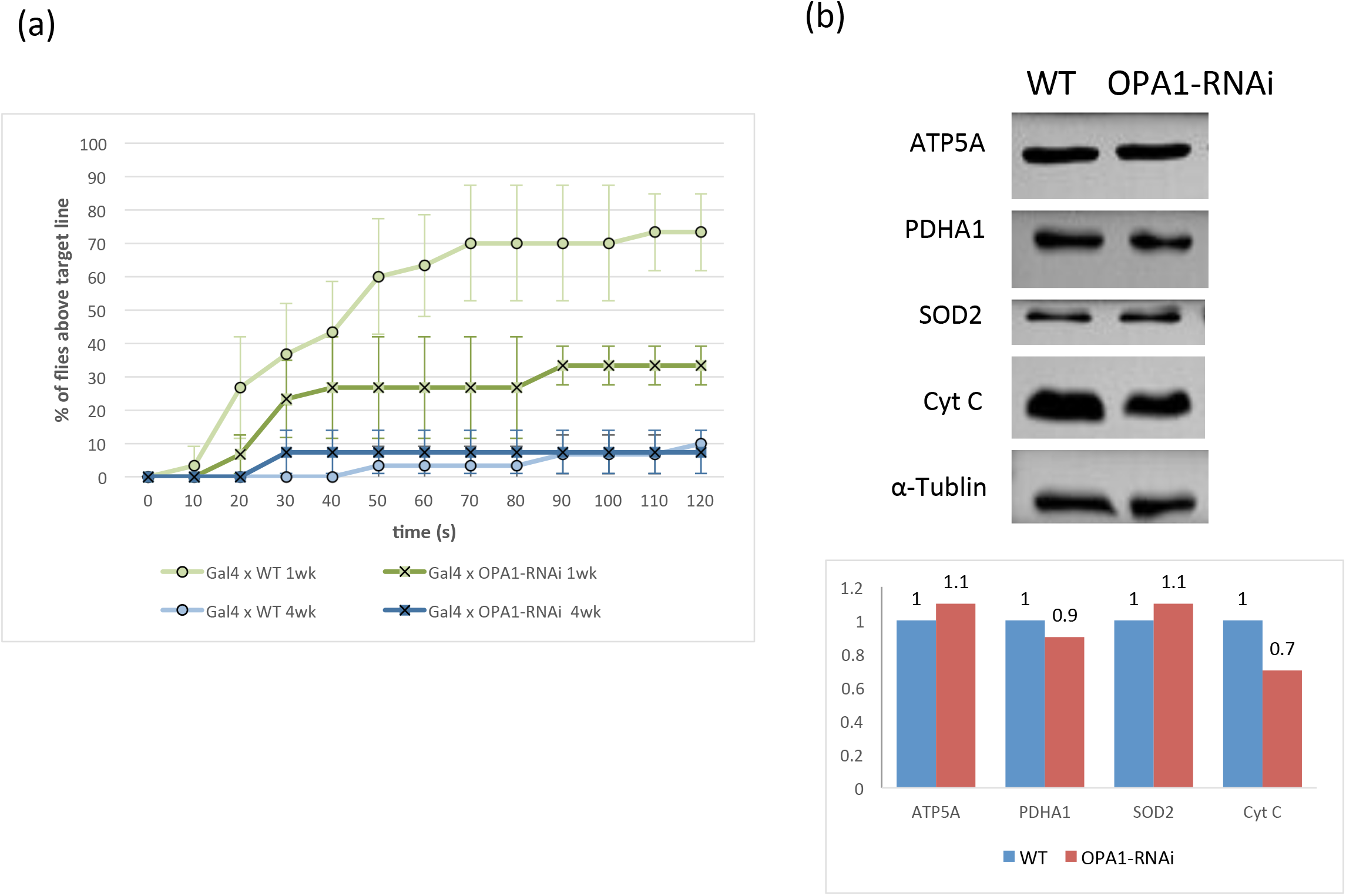
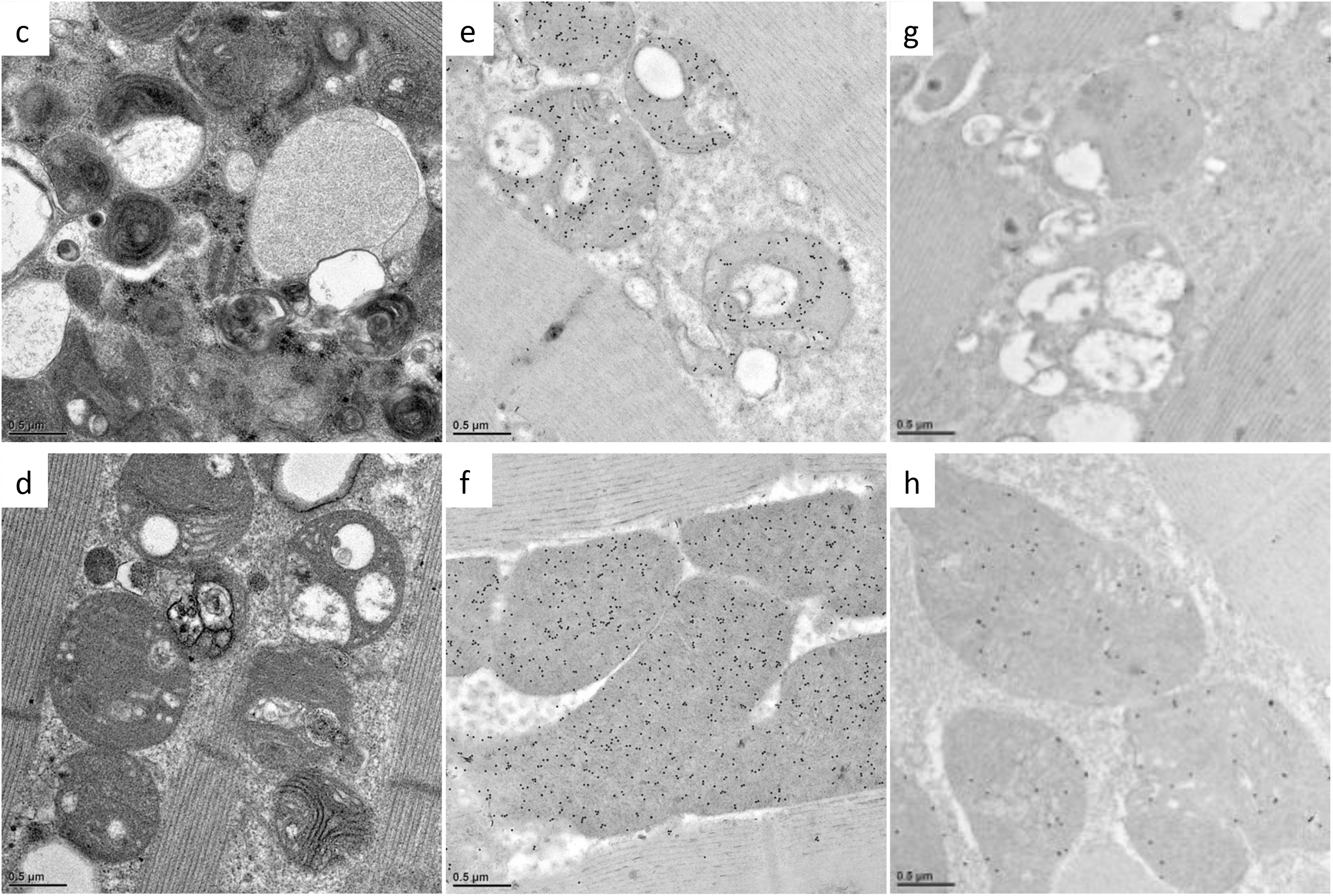

**Fig S4.**
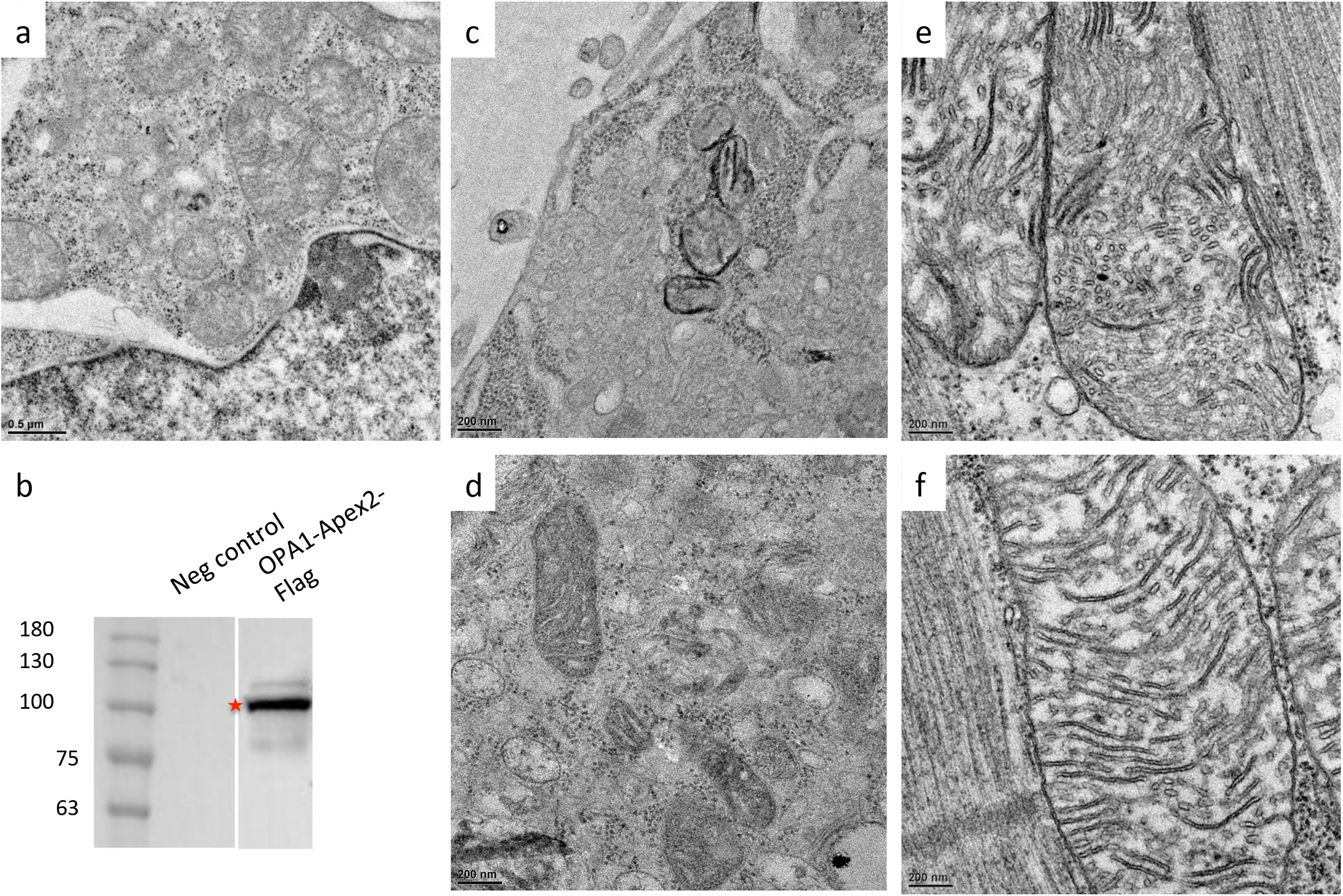

