## Supplemental material for "Mitochondrial cristae biogenesis coordinates with ETC complex IV assembly during *Drosophila* maturation"

**Fig S1. Characterization of the knock-in flies containing Apex2-conjugation**

(a) COX4-Apex2-Flag, OXA1-Apex2-Flag, and OPA1-Apex2-Flag expression of the knock-in flies was detected by western blot using anti-flag antibody. (b) Illustration of the mitochondrial location of c-terminal COX4-Apex2 tag. (c) Western blot analysis of S2 cells overexpressing *D. melanogaster* OXA1-Apex2-Flag using anti-flag antibody. Red stars indicate predicted protein-Apex2 conjugates.

**Fig S2. Characterization of Marf-knockdown flies**

(a) The climbing assay was performed on Marf-knockdown flies at day 1 and week 4 and compared with the wild-type cross. (b) Western blot analysis and quantification of mitochondrial proteins, including ATP5A, PDHA1, SOD2, and Cyt c, in the whole fly extracts of the wild type and Marf-knockdown at week 4. The relative protein abundance was quantified by the densitometry and normalized to the signal of α-tubulin. The ratios were subsequently normalized to those of the wild type at week 4. Marf-knockdown flies at week 1 were analyzed by thin-section TEM (c) and COX activity staining (d). Marf-knockdown flies at week 4 were analyzed by immuno-EM against Cyt c (f). The wild type controls for immuno-EM against ATP5A and Cyt c were shown in (e) and (g), respectively. (h) Illustration of the mitochondrial location of the c-terminal ATP synthase OSCP-Apex2 tag.

**Fig S3. Characterization of OPA1-knockdown flies**

(a) The climbing assay was performed on OPA1-knockdown flies at day 1 and week 4 and compared with the wild-type cross. (b) Western blot analysis and the quantification of mitochondrial proteins, including ATP5A, PDHA1, SOD2, and Cyt c in the whole fly extracts of the wild type and OPA1-knockdown at week 4. The relative protein abundance was quantified by the densitometry and normalized to the signal of α-tubulin. The ratios were subsequently normalized to those of the wild type. OPA1-knockdown flies at week 4 were analyzed by thin-section TEM (c), COX activity staining (d), and immuno-EM against ATP5A (e) and Cyt c (g). The wild-type controls for immuno-EM are shown in (f) and (h), respectively.

**Fig S4. Ultrastructural tracking of OPA1**

(a) Negative control of Apex2 staining of mock-transfected 293T cells. (b) Western blot analysis of 293T overexpressing OPA1-Apex2 using anti-flag antibody. (c) Apex2 staining of S2 cells expressing OPA1-Apex2-Flag and (d) the mock-transfection. (e) Apex2 staining of the IFM of OPA1-Apex2 knock-in flies at day 1 and (f) the wild type at day 1 as the negative control.

**Movie 1. Serial-section electron tomography and volume segmentation of *Drosophila* upon eclosion.**

**Movie 2. Serial-section electron tomography and volume segmentation of *Drosophila* upon eclosion stained for COX activity.**

**Movie 3. Serial-section electron tomography and volume segmentation of Marf-knockdown fly at week 4.**

**Movie 4. Serial-section electron tomography of OPA1-knockdown fly at week 4 stained for COX activity.**
